# PEhub resolves the hierarchical regulatory architecture of multi-way enhancer hubs in the human brain

**DOI:** 10.64898/2026.02.09.704931

**Authors:** Jiang Tan, Yidan Sun

## Abstract

Chromatin interaction assays capture regulatory architecture as stochastic pairwise contacts, limiting the ability to resolve how multiple enhancers cooperatively regulate transcription. Here we introduce a promoter-centric quantitative framework, termed PEhub, that resolves multi-way enhancer hubs as higher-order regulatory units from chromatin interaction data. By reparameterizing stochastic pairwise ligation events into promoter-conditioned enhancer networks, our approach explicitly models synergistic enhancer cooperation while accounting for distance-dependent interaction decay through a statistically principled null model. Using H3K27ac HiChIP data, we identify promoter-anchored enhancer hubs and validate their physical existence with single-molecule Pore-C, demonstrating that inferred hubs correspond to bona fide multi-way chromatin assemblies. Application to six human brain regions reveals that enhancer hubs are associated with elevated transcriptional output and exhibit a hierarchical organization spanning shared, circuit-specific, and region-restricted regulatory programs. This architecture hierarchically stratifies genetic risk and transcription factor deployment, linking three-dimensional genome organization to transcriptional control and disease-associated variation. Together, this promoter-centric framework provides a generalizable strategy for resolving higher-order regulatory architecture from 3D genome data and establishes multi-way enhancer hubs as a functionally and genetically meaningful layer of transcriptional regulation in complex tissues.

## Introduction

Enhancers play a central role in orchestrating precise spatiotemporal gene regulation, particularly in complex tissues such as the human brain^1–5^. For decades, enhancers were studied individually, each viewed as an independent regulatory switch^6,7^. However, emerging evidence shows enhancers frequently act in cooperative groups, working together to fine tune transcriptional output and confer robustness to gene regulation^8^. In this context, multiple enhancers can converge on a shared promoter to form higher-order regulatory assemblies, often referred to as multi-way enhancer hubs^3,9^ (Fig.1). Their composition and activity are highly dynamic, some enhancer hubs are conserved across cell types, while others are unique to specialized state^10^. Disruption of these cooperative enhancer networks, rather than misregulation of individual enhancer–promoter pairs, can rewire transcriptional programs, driving various diseases^11–13^.

**Figure 1:**
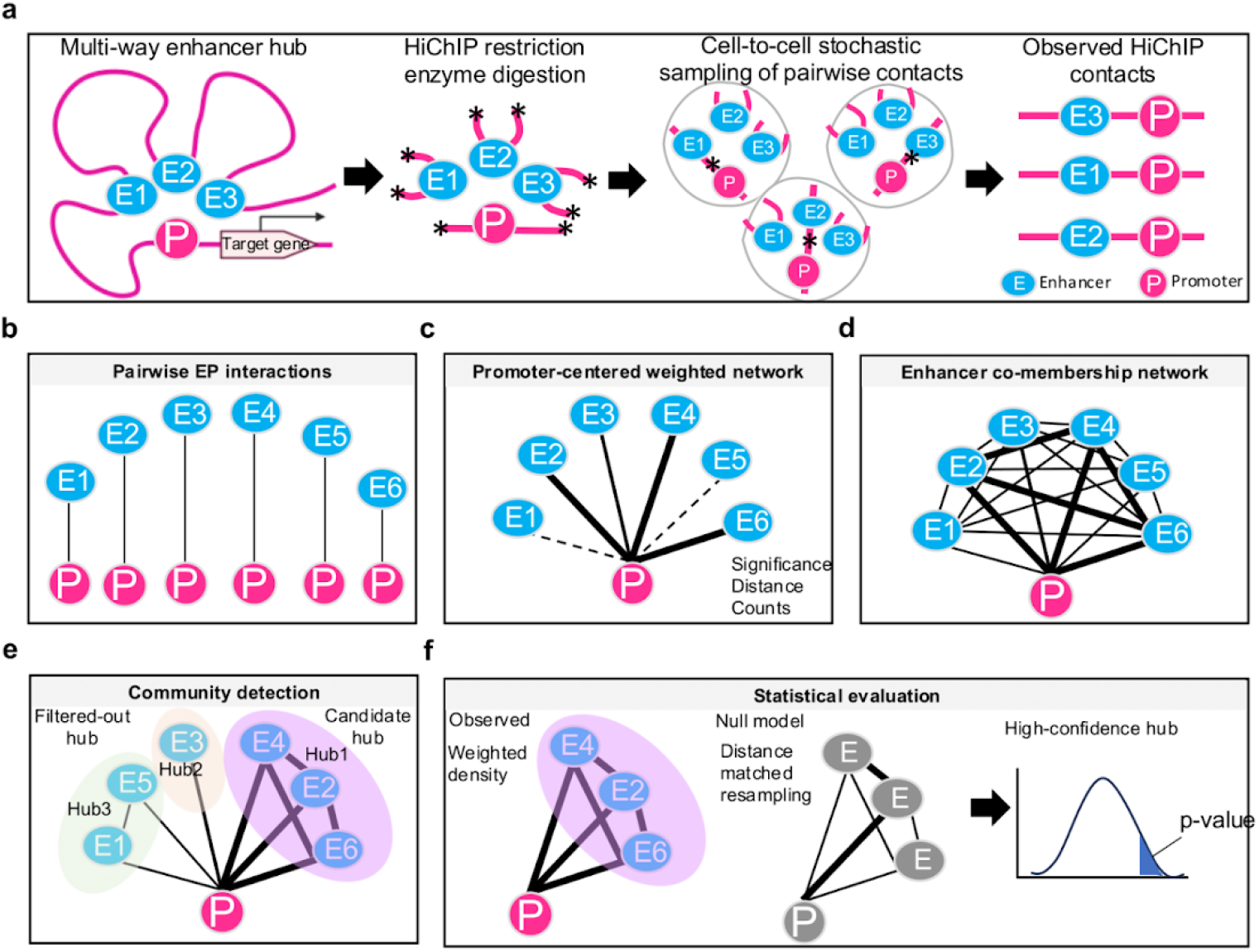
A promoter-centric quantitative framework defines multi-way enhancer hubs as high-order regulatory units. **a**, Conceptual illustration of how underlying multi-way enhancer–promoter assemblies are observed as pairwise contacts in HiChIP data. In the nucleus, a promoter can be simultaneously co-localized with multiple enhancers within a higher-order chromatin complex. During HiChIP, proximity ligation captures only a subset of pairwise contacts from this multi-way assembly, and different cells stochastically sample different enhancer–promoter pairs. Subsequent fragmentation and short-read sequencing further collapse higher-order organization into independent pairwise enhancer–promoter interactions. **b**, Pairwise enhancer–promoter (EP) interactions detected by HiChIP, which constitute the primary input to the framework. **c**, Promoter-centric reparameterization of pairwise interactions. For each promoter, enhancer–promoter interactions are modeled as a local weighted network, in which enhancer–promoter edges are assigned quantitative weights that integrate interaction strength, statistical confidence and distance dependency. **d**, Construction of promoter-conditioned enhancer co-membership networks. For each promoter, enhancer–enhancer co-membership scores quantify the extent of cooperative contribution between enhancers conditional on their shared promoter, yielding a weighted enhancer–enhancer graph. **e**, Identification of multi-way enhancer hubs by community detection. Leiden clustering is applied to each promoter-centered co-membership network, allowing multiple enhancer hubs per promoter. Hubs are ranked by dominance and the highest-ranking hubs per promoter were retained as candidate hubs and the rest are termed as filtered-out hubs. **f**, Statistical evaluation of enhancer hubs using a distance-stratified null model. Observed hub scores (weighted density) are compared against null distributions generated by distance-matched resampling of enhancer–promoter interaction weights, enabling empirical p-value estimation and identification of high-confidence multi-way enhancer hubs.

Despite the recognition of its biological importance, current analytical paradigms lack a formal operational definition for resolving multi-way enhancer hubs as functional transcriptional units, leaving a critical gap between higher-order chromatin topology and downstream gene regulation. Existing methods typically rely on post hoc aggregation of pairwise loops into global interaction graphs, followed by spectral clustering^14^ or inflection-point–based^15^ thresholds to partition communities. In the absence of explicit correction for the intrinsic distance-dependent decay of chromatin interactions, such formulations can preferentially capture proximal elements, confounding spatial proximity with functional coordination. Moreover critically, by modeling regulatory architecture as a flat network of edges, these approaches do not explicitly capture synergistic enhancer–enhancer cooperation, nor do they recognize the promoter as a functional integrator of multi-way regulatory input. Without a statistically principled null model to distinguish coherent synergistic assemblies from stochastic spatial proximities, it remains unclear whether identified “hubs” correspond to bona fide multi-way physical structures or arise from local interaction density and coverage effects. Consequently, the field lacks a generalizable framework for defining enhancer hubs as promoter-anchored transcriptional regulatory units, thereby precluding systematic linkage of higher-order regulatory organization to transcriptional output.

Here, we introduce a promoter-centric, weighted network framework for identifying multi-way enhancer hubs as high-order regulatory units from chromatin interaction (e.g., H3K27ac HiChIP^16,17^) data. Rather than aggregating pairwise interactions at the genomic region level, our approach models enhancer–promoter interactions as local weighted networks anchored at individual promoters. By constructing a normalized co-membership matrix, we explicitly quantify the synergistic co-occurrence of multiple enhancers, prioritizing those that act as coherent functional units. A critical innovation of our framework is the implementation of a distance-stratified null model, which enables the computation of empirical p-values and effect sizes, thereby providing a robust statistical foundation for hub identification. This framework enables quantitative characterization of hub architecture, including hub size, internal coherence, dominance, and stability, and provides a natural interface for integrating transcriptional and epigenomic measurements. Most importantly, we leverage long-read Pore-C^18^ data to provide direct experimental evidence that our computationally inferred hubs correspond to physical multi-way chromatin assemblies, bridging the gap between statistical inference and molecular architecture.

Applying this framework to six human brain regions, we demonstrate that multi-way enhancer hubs are robustly associated with transcriptional output, providing quantitative evidence that higher-order regulatory assemblies contribute to gene regulation in complex tissues. More broadly, this framework enables the systematic comparison of enhancer hub architecture across regions, revealing structured patterns of hub sharing and specialization that reflect distinct regulatory programs. These analyses illustrate how promoter-centric hub definitions can be leveraged to interrogate tissue- and context-specific regulatory organization at scale. Together, this work establishes a generalizable framework for resolving high-order regulatory architecture from 3D genome data. By integrating promoter-centric hub inference, long-read validation, and cross-brain-region analysis, our approach bridges multi-way chromatin organization with transcriptional control and disease-associated variation, providing a foundation for systematic interrogation of complex regulatory systems in development and disease.

## Results

### A promoter-centric quantitative framework defines multi-way enhancer hubs as high-order regulatory units

While standard HiChIP captures chromatin architecture as a collection of pairwise contacts that are stochastically sampled from underlying multi-way chromatin assemblies (e.g., Enhancer1–Promoter, Enhancer2–Promoter), it does not directly resolve the coordinated activity of multiple enhancers acting on a shared promoter (Fig. 1a). To formally resolve multi-way enhancer hubs as transcriptional regulatory units, we developed a promoter-centric framework, termed *PEhub*, that reparameterize these pairwise fragments (Fig. 1b) into a higher-order network effectively reconstructing the synergistic multi-enhancer assemblies that are fragmented during conventional sequencing library preparation. Consequently, our framework exposes a layer of cis-regulatory organization that is inaccessible to loop-centric formulations.

In this framework, each promoter is treated as an explicit regulatory reference point that defines a local interaction neighborhood (Fig. 1c). Instead of analyzing enhancer-promoter interactions globally, we reconstruct the regulatory landscape of each promoter as a network of connected enhancers, represented as nodes whose edges reflect their regulatory influence on the promoter. This promoter-centric reparameterization aligns the analysis directly with transcriptional output and provides a natural scaffold for comparing regulatory architectures across genes and biological contexts.

A central design principle of this framework is the explicit modeling of relative enhancer contribution (Fig. 1c). Rather than relying on binary interaction calls, enhancer–promoter interactions are represented as weighted edges that integrate interaction strength, statistical confidence, and genomic distance. This distance-aware formulation accounts for interaction decay and technical heterogeneity, enabling robust comparison of enhancer contributions both within and across promoters while avoiding overemphasis of proximal or highly covered interactions.

To identify coordinated enhancer activity, we quantify enhancer co-membership relationships that assess whether pairs of enhancers jointly contribute to promoter regulation beyond their individual effects (Fig. 1d). These co-membership relationships transform promoter-centered interaction neighborhoods into structured enhancer–enhancer networks, in which patterns of cooperation emerge naturally. Community detection is then applied to these networks to identify enhancer hubs as modular, densely interconnected assemblies without requiring prior assumptions about hub size, enhancer number, or genomic span (Fig. 1e). Importantly, this formulation allows multiple enhancer hubs to coexist at a single promoter, reflecting the modular and context-dependent nature of cis-regulatory programs.

To ensure the identified hubs represent bona fide regulatory assemblies rather than stochastic spatial proximities, we implemented a distance-matched null model (Fig. 1f). By resampling enhancer weights from global, distance-stratified pools, we generated a background distribution that preserves the data’s original distance-dependency and interaction heterogeneity while disrupting coordinated structures. This allows for the computation of empirical p-values and effect sizes for each hub, providing a principled means to discriminate genuine multi-enhancer organization from artifacts of interaction density or genomic coverage. The robustness of hub structure is further assessed within this framework through perturbation-based resampling, enabling a principled means to evaluate sensitivity to sampling and clustering variability.

Together, this promoter-centric framework transforms pairwise chromatin interaction data into a higher-order regulatory representation that explicitly models enhancer cooperation. By integrating promoter anchoring, quantitative weighting, co-membership–based network inference, and statistical evaluation, it enables systematic identification, quantification, and comparison of enhancer hubs across biological contexts. These results establish multi-way enhancer hubs as a distinct and functionally meaningful layer of cis-regulatory organization, bridging chromatin architecture with downstream transcriptional control in complex contexts.

### Direct physical validation of multi-way enhancer hubs using Pore-C

To test whether the multi-way enhancer hubs inferred from our framework correspond to physical higher-order chromatin assemblies, we leveraged long-read Pore-C data from GM12878 cells^18^. Unlike conventional Hi-C, Pore-C directly captures multi-contact chromatin interactions within individual DNA concatemers, providing a “ground truth” for higher-order architecture^18^. We defined a Pore-C read as hub-supporting if it concurrently overlapped at least three hub elements, including enhancers and/or the associated promoter (total contact order k≥3) (Fig. 2a; Supplementary Fig. 1).

**Figure 2:**
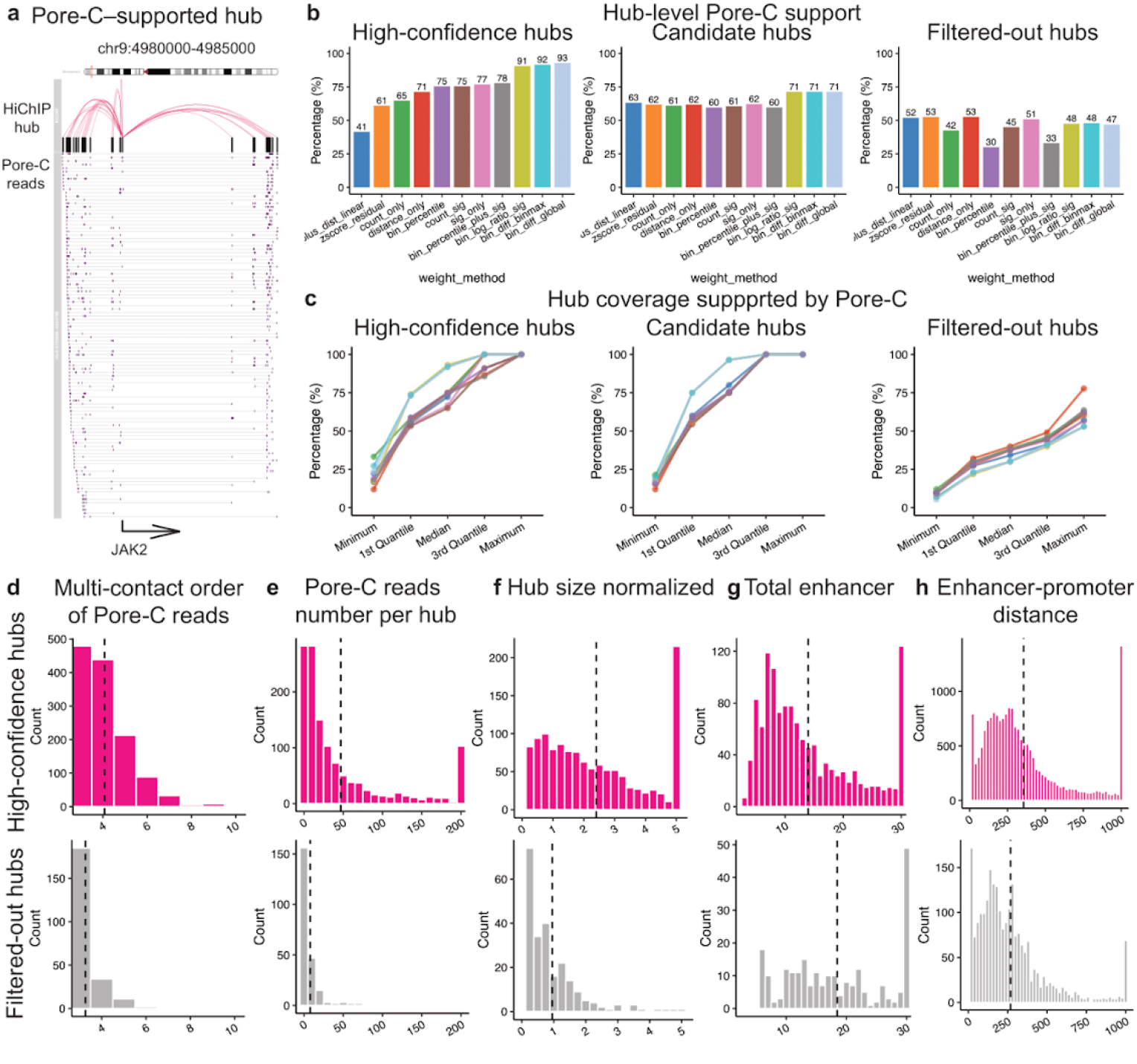
Direct physical validation of multi-way enhancer hubs using Pore-C. **a**, A representative example of high-confidence multi-way enhancer hubs identified from GM12878 H3K27ac HiChIP data and independently supported by single-molecule Pore-C reads. A hub is considered supported if individual Pore-C reads overlap at least three hub elements, including enhancers and/or the associated promoter. For each locus, HiChIP-inferred enhancer hub structures (top) are shown together with aligned Pore-C multi-contact reads (bottom), illustrating direct physical evidence for higher-order chromatin assemblies. **b**, Fraction of inferred enhancer hubs supported by multi-contact Pore-C reads across eleven interaction weighting strategies. Candidate hubs identified by community detection and high-confidence hubs retained after statistical significance and stability filtering are compared with filtered-out hubs. **c**, Distribution of hub coverage, defined as the fraction of enhancers within each hub supported by Pore-C reads, across weighting strategies and hub categories. **d**, Distribution of multi-contact order (k) for Pore-C reads supporting enhancer hubs, reflecting higher-order chromatin interactions beyond pairwise contacts. High-confidence hubs are compared with filtered-out hubs. **e**, Distribution of the total number of supporting Pore-C reads per hub for high-confidence and filtered-out hubs. **f**, Supporting Pore-C reads per hub normalized by the number of enhancers within each hub, controlling for hub size effects. **g**, Distribution of hub size for high-confidence and filtered-out hubs. **h**, Genomic distances between enhancers and promoters within hubs, demonstrating that increased Pore-C support is not attributable to increased spatial proximity.

We first quantified the fraction of inferred hubs supported by multi-contact Pore-C reads (Fig. 2b). Across multiple interaction weighting strategies, candidate enhancer hubs identified by community detection exhibited consistent physical support, with approximately 60–70% of hubs supported by Pore-C reads. In contrast, filtered-out hubs showed substantially lower support (30–53%). Notably, after further selection based on statistical significance and stability (FDR<0.05 & stability > 0.5, see Methods for details), three weighting schemes, including *bin_sig_ratio, bin_diff_max*, and *bin_diff_global*, yielded high-confidence hubs with Pore-C support exceeding 90%, indicating strong enrichment for physically coherent multi-way structures.

To assess whether Pore-C support was broadly distributed across hub components instead of confined to a small subset of enhancers, we next examined hub coverage, defined as the fraction of enhancers within each hub that were supported by Pore-C hub-supporting reads (Fig. 2c). Across most weighting strategies, median high-confidence hub coverage ranged from 55% to 75%, whereas the *bin_sig_ratio* and *bin_diff_max* methods achieved median coverage values approaching 90%. In contrast, filtered-out hubs displayed markedly lower coverage, with median values below 40%, indicating that Pore-C support is both more frequent and more uniformly distributed within high-confidence hubs.

Based on their consistently high Pore-C support and coverage, we selected the *bin_sig_ratio* weighting strategy for subsequent analyses. To determine whether Pore-C validation reflected genuine multi-way chromatin organization rather than accumulation of independent pairwise interactions, we examined the distribution of multi-contact order (*k*) for Pore-C reads associated with inferred hubs (Fig. 2d). High-confidence hubs exhibited higher median *k* values than filtered-out hubs, consistent with enrichment for higher-order chromatin interactions.

To quantify the strength of Pore-C evidence, we compared the distribution of supporting multi-contact reads per hub. High-confidence hubs showed increased Pore-C hub-supporting reads relative to filtered-out hubs (Fig. 2e). To exclude the possibility that increased Pore-C reads were driven by hub size, we normalized the number of supporting reads by the number of enhancers within each hub (Fig. 2f). Even after normalization, high-confidence hubs showed greater median Pore-C support reads than filtered-out hubs, indicating that enrichment is not attributable to trivial scaling effects.

Finally, we examined hub size and enhancer–promoter distance distributions to assess whether physical validation was confounded by simple structural properties. High-confidence hubs were, on average, smaller than filtered-out hubs (Fig. 2g), while exhibiting more genomically dispersed (Fig. 2h), arguing against hub compactness or physical proximity as explanations for increased Pore-C support.

Results were further compared to other weighting strategies for all Pore-C validation metrics (Supplementary Figs. 2-3).

Together, these analyses provide independent, orthogonal physical validation that multi-way enhancer hubs identified by our framework correspond to bona fide higher-order chromatin assemblies. The observed enrichment for multi-contact interactions, uniform internal support, and resistance to size and distance confounders indicates that these hubs capture coordinated multi-enhancer regulatory organization that cannot be explained by independent pairwise interactions or simple structural confounders.

### Enhancer hubs are associated with elevated transcriptional output across human brain regions

We next applied our framework to H3K27ac HiChIP data from six human brain regions, including substantia nigra, middle frontal gyrus, caudate, hippocampus, parietal lobe and superior temporal gyrus, each profiled with two biological replicates^1,5^. To ensure robustness, we defined replicable enhancer hubs as candidate hubs identified at the same promoter in both replicates, sharing at least two enhancers, and significant in at least one replicate (FDR ≤ 0.05). Across regions, we identified markedly different numbers of reproducible enhancer hubs, ranging from 1,460 in substantia nigra and 1,362 in middle frontal gyrus to substantially fewer hubs in parietal lobe and superior temporal gyrus (Fig. 3a; Supplementary Table 1). These differences likely reflect a combination of biological heterogeneity and region-specific regulatory complexity, rather than technical artifacts, as identical analytical criteria were applied to all datasets.

**Figure 3:**
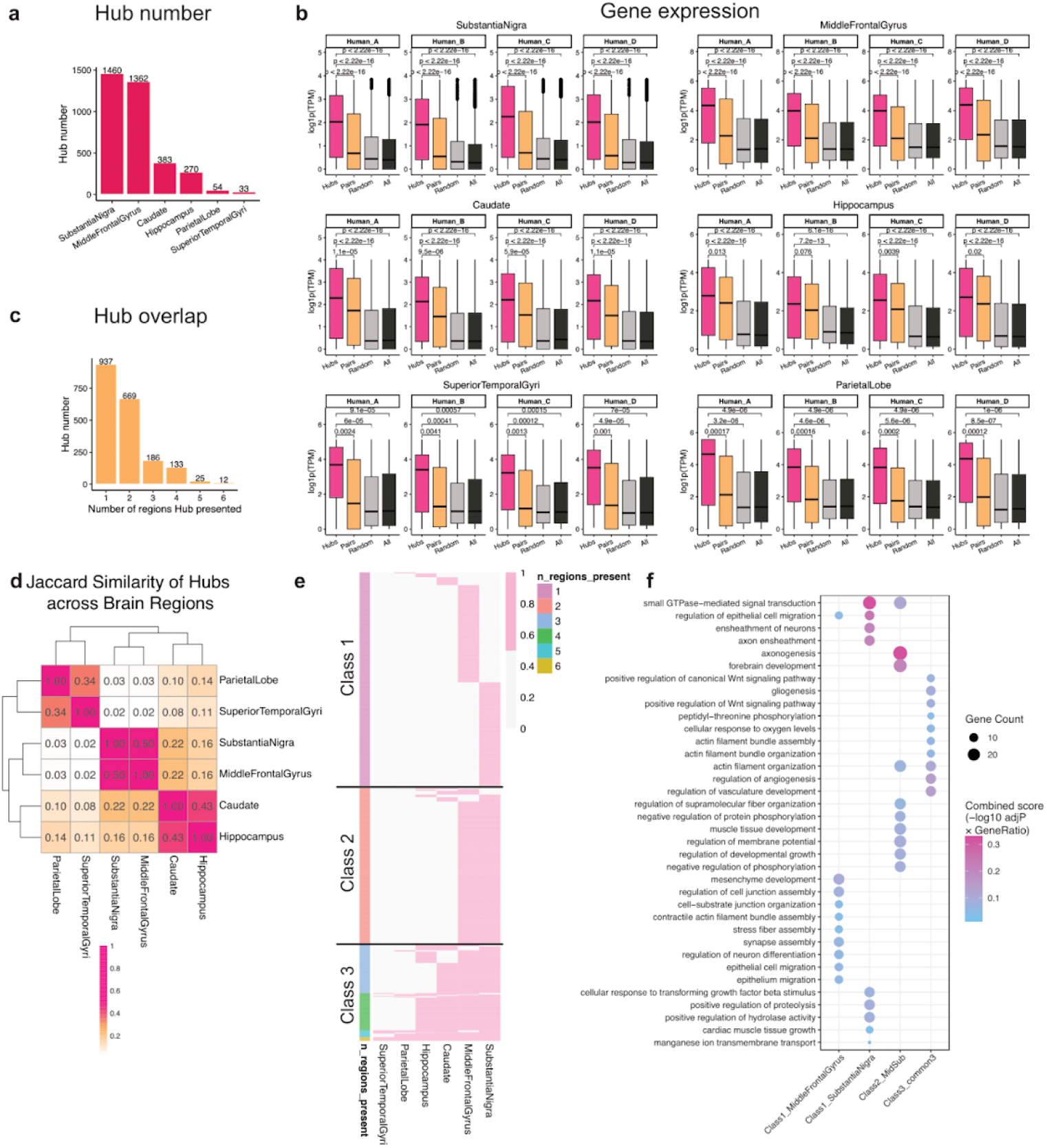
Hierarchical organization of multi-way enhancer hubs across human brain regions. **a**, Number of replicable multi-way enhancer hubs identified across six human brain regions (substantia nigra, middle frontal gyrus, caudate, hippocampus, parietal lobe and superior temporal gyrus). Each region was profiled with two biological replicates. Replicable enhancer hubs were defined as hubs identified at the same promoter in both replicates, sharing at least two enhancers, and significant in at least one replicate. **b**, RNA-seq expression levels (TPM) of genes associated with enhancer hubs, pairwise enhancer–promoter interactions, randomly selected genes (n = 1,000), and all other expressed genes, shown separately for each brain region and individual. Hub-associated genes consistently exhibit higher expression relative to all control gene sets. Boxplots indicate median and interquartile range; whiskers denote 1.5× interquartile range. **c**, Distribution of hub-associated genes stratified by the number of brain regions in which they are associated with enhancer hubs, highlighting region-specific and shared regulatory programs. **d**, Pairwise overlap of hub-associated gene sets across brain regions, quantified by the Jaccard index, revealing structured sharing between anatomically and functionally related regions. **e**, Heatmap summarizing overlap of hub-associated gene sets between brain regions, highlighting both region-specific regulatory programs and subsets of genes shared across related regions. Based on regional sharing patterns, enhancer hubs were classified into three hierarchical regulatory classes: class 1, region-restricted hubs observed in a single brain region; class 2, pair-specific hubs shared exclusively between two regions; and class 3, multi-region shared hubs observed across three or more brain regions. **f**, Gene Ontology enrichment analysis of hub-associated genes stratified by regional sharing class, highlighting conserved neuronal processes among shared hubs, pair-specific and region-biased biological programs among region-specific hubs.

To evaluate the functional relevance of enhancer hubs, we compared RNA-seq expression levels of hub-associated genes with several control gene sets, including genes linked to individual enhancer–promoter interactions, randomly sampled genes, and all other expressed genes within each region. Across all six brain regions, genes associated with enhancer hubs exhibited significantly higher expression than all control groups (Fig. 3b). This observation is consistent with prior studies implicating enhancer hubs and multi-enhancer configurations in robust transcriptional activation^3,8,9^, and supports the notion that coordinated multi-enhancer assemblies represent a distinct and potent mode of cis-regulatory control in the human brain.

### Hierarchical organization of multi-way enhancer hubs across human brain regions

To assess the extent of regulatory sharing across brain regions, we quantified the number of regions in which each hub-associated gene was observed. This analysis revealed a highly non-uniform distribution of regulatory sharing (Fig. 3c), delineating three dominant sharing patterns: genes shared across three or more regions (class 3, 18%), genes shared between exactly two regions (class 2, 34%), and genes restricted to a single region (class 1, 48%).

To determine whether pair-wise sharing reflected structured inter-regional relationships rather than stochastic overlap, we quantified overlap of hub-associated genes using the Jaccard index (Fig. 3d). This analysis revealed pronounced structure, with strong pair-wise overlap observed between substantia nigra and middle frontal gyrus, between caudate and hippocampus, and between parietal lobe and superior temporal gyrus. These region pairs correspond to known functional and anatomical relationships, suggesting that pair-wise enhancer hub sharing reflects coordinated regulatory programs rather than stochastic overlap.

We next examined whether pair-wise shared genes represented discrete modules or subsets of broader shared programs. Notably, whereas most pair-wise overlaps were largely attributable to genes also shared across multiple regions, the middle frontal gyrus–substantia nigra pair emerged as a distinct exception. Specifically, 602 hub-associated genes were shared exclusively between these two regions and were not observed in enhancer hubs from other regions, identifying a discrete pair-specific enhancer hub module linking middle frontal gyrus and substantia nigra (Fig. 3e).

In addition to shared regulation, nearly half of hub-associated genes (48%) were restricted to a single brain region. Region-specific enhancer hub regulation was unevenly distributed across the brain: substantia nigra and middle frontal gyrus exhibited the highest degree of specificity, with approximately 30% of their hub-associated genes restricted to their respective regions (403/1,362 in middle frontal gyrus and 485/1,460 in substantia nigra). In contrast, region-specific genes accounted for less than 10% of hub-associated genes in caudate, hippocampus, parietal lobe and superior temporal gyrus (Fig. 3e).

To elucidate the biological meaning of these sharing patterns, we performed Gene Ontology enrichment analyses on hub-associated genes stratified by their regional specificity (Fig. 3f). Genes shared across multiple brain regions were enriched for conserved signaling and regulatory programs, including Wnt signaling, phosphorylation-associated regulation, vascular and oxygen-responsive pathways, and cytoskeletal organization, defining a shared regulatory backbone deployed across brain regions. In contrast, the middle frontal gyrus–substantia nigra pair-specific module showed selective enrichment for axonogenesis and forebrain development, together with coordinated regulation of signaling, growth, and membrane excitability, consistent with a structured morphogenetic program underlying circuit-level organization.

At the most specific level, region-restricted enhancer hubs exhibited distinct functional biases (Fig. 3f). Substantia nigra–specific hubs were enriched for neuronal and axonal ensheathment together with small GTPase–mediated signal transduction, growth factor–responsive signaling (cellular response to transforming growth factor beta stimulus), and proteolysis/hydrolase regulation, indicating emphasis on structural support and remodeling programs that may underlie long-range organization and selective vulnerability. Middle frontal gyrus–specific hubs preferentially targeted regulators of cell junction assembly and cell–substrate junction organization, cytoskeletal contractility (stress fiber and contractile actin bundle assembly), synapse assembly, and neuronal differentiation, consistent with region-biased control of connectivity and differentiation state.

Together, these results reveal a hierarchical regulatory architecture of multi-way enhancer hubs in the human brain. Multi-region shared hubs establish a conserved neuronal signaling backbone (class 3), discrete pair-specific modules encode coordinated circuit-level morphogenetic programs (class 2), and region-restricted hubs bias regulatory activity toward region-specific functions (class 1). This layered organization illustrates how the promoter-centric framework enables systematic dissection of higher-order regulatory organization across complex tissues and provides a principled basis for understanding how shared neuronal programs are integrated with region- and circuit-specific regulatory control.

### GWAS stratification reveals hierarchical genetic risk embedded within enhancer hub architecture

To examine whether disease-associated genetic variation is distributed across higher-order regulatory architecture, we analyzed the overlap between enhancer hubs and genome-wide association study (GWAS) variants^19^. A total of 713,013 unique single-nucleotide polymorphisms (SNPs) were compiled from 31,944 GWAS traits cataloged in the UCSC Genome Browser’s GWAS track and quantified their intersection with enhancer hub elements identified across six human brain regions.

Across all three enhancer hub classes, GWAS variants spanned a broad range of phenotypic categories, including molecular traits, physiological measures and brain-related phenotypes (Fig. 4a), reflecting the pleiotropic nature of regulatory variation. Given our focus on brain regulatory architecture, we next concentrated on brain-relevant traits, where clear and hierarchical patterns of genetic risk emerged.

**Figure 4:**
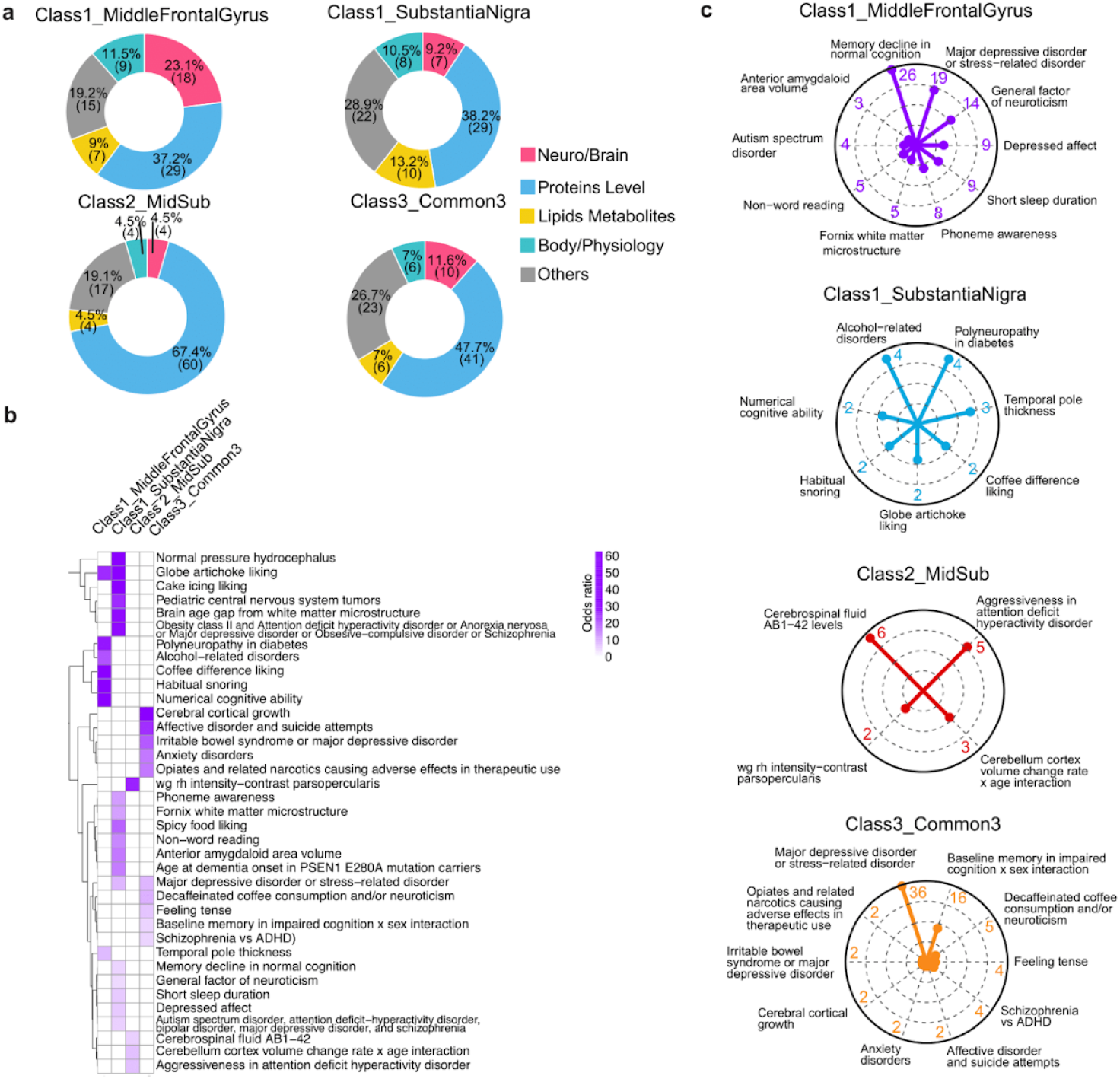
GWAS stratification reveals hierarchical genetic risk embedded within enhancer hub architecture. **a**, Distribution of GWAS trait categories represented among variants overlapping enhancer hubs across six human brain regions, stratified by enhancer hub class. Enhancer hub classes are defined by regional sharing of hub-associated genes: Class 1, region-restricted hubs observed in a single brain region; Class 2, pair-specific hubs shared exclusively between two regions; and Class 3, multi-region shared hubs observed in three or more regions. Trait categories span neuro/brain-related phenotypes, protein-level traits, lipid and metabolite traits, physiological measures, and other complex traits, reflecting the pleiotropic nature of regulatory variation. **b**, Enrichment of brain-related GWAS traits across enhancer hub classes. Odds ratios are shown for representative traits enriched within region-restricted hubs (Class 1), pair-specific hubs linking middle frontal gyrus and substantia nigra (Class 2), and multi-region shared hubs (Class 3). Only traits with enrichment p-value ≤ 0.01 and odds ratio ≥ 5 are displayed. **c**, Radar plots summarizing the SNP number of enriched brain-related GWAS traits across Class 1, Class 2, and Class 3 enhancer hubs, corresponding to the traits shown in **b**. Together, these analyses indicate that genetic risk for brain-related traits is hierarchically organized across enhancer hub classes, with progressively increased specialization from multi-region shared to region-restricted regulatory architectures.

Variants linked to broadly shared affective and neuropsychiatric traits, including anxiety-related phenotypes, schizophrenia and suicide-associated traits, were preferentially enriched within multi-region shared enhancer hubs (Class 3) (Fig. 4b,c). This pattern is consistent with perturbation of conserved neuronal regulatory programs that likely underlie generalized neuropsychiatric vulnerability.

In contrast, pair-specific enhancer hubs linking the middle frontal gyrus and substantia nigra (Class 2) showed selective enrichment for traits related to brain structural dynamics and circuit-level modulation, including cerebrospinal fluid Aβ1–42 levels, age-dependent cortical volume changes and impulsivity-related behavioral traits (Fig. 4b,c). These results indicate that a substantial fraction of genetic risk is concentrated within discrete circuit-level regulatory modules that are superimposed on a shared neuronal regulatory backbone.

At the most specialized level, we observed a pronounced concentration of cognitive and neuropsychiatric risk within region-restricted enhancer hub modules (Class 1). Middle frontal gyrus–specific hubs were uniquely enriched for variants linked to high-level cognitive traits, such as phoneme awareness and non-word reading, alongside complex neurodevelopmental and psychiatric conditions (Fig. 4b,c). Notably, these hubs also captured genetic associations with late-life cognitive trajectories, suggesting that region-specific higher-order regulatory programs may influence both cognitive development and progressive vulnerability across the lifespan. In contrast, substantia nigra–specific hubs prioritized a distinct phenotypic spectrum, exhibiting enrichment for subcortical, metabolic and autonomic-related traits, including alcohol-related disorders and habitual snoring, consistent with the region’s specialized roles in reward processing and homeostatic regulation (Fig. 4b,c).

Collectively, these findings demonstrate that genetic risk for brain-related traits is not uniformly distributed across regulatory space but is stratified across hierarchical layers of enhancer hub architecture. Multi-region shared enhancer hubs capture broadly conserved neuronal risk, whereas pair-specific and region-restricted hubs progressively concentrate genetic risk within increasingly specialized circuit- and region-level regulatory programs. This hierarchical organization provides a principled framework for interpreting non-coding disease-associated variation in the context of higher-order regulatory architecture.

### Transcription factor programs underlying hierarchical enhancer hub regulation

To characterize the transcriptional programs driving hierarchical enhancer hub organization, we systematically analyzed transcription factor (TF) binding patterns within multi-way enhancer hubs across human brain regions using scPrinter^20^, an AI-based footprinting framework applied to matched ATAC-seq data^1,5^ (Fig. 5a). This approach enables bin-level inference of TF occupancy across enhancer hub elements, providing a means to interrogate regulatory programs operating within higher-order chromatin assemblies.

**Figure 5.**
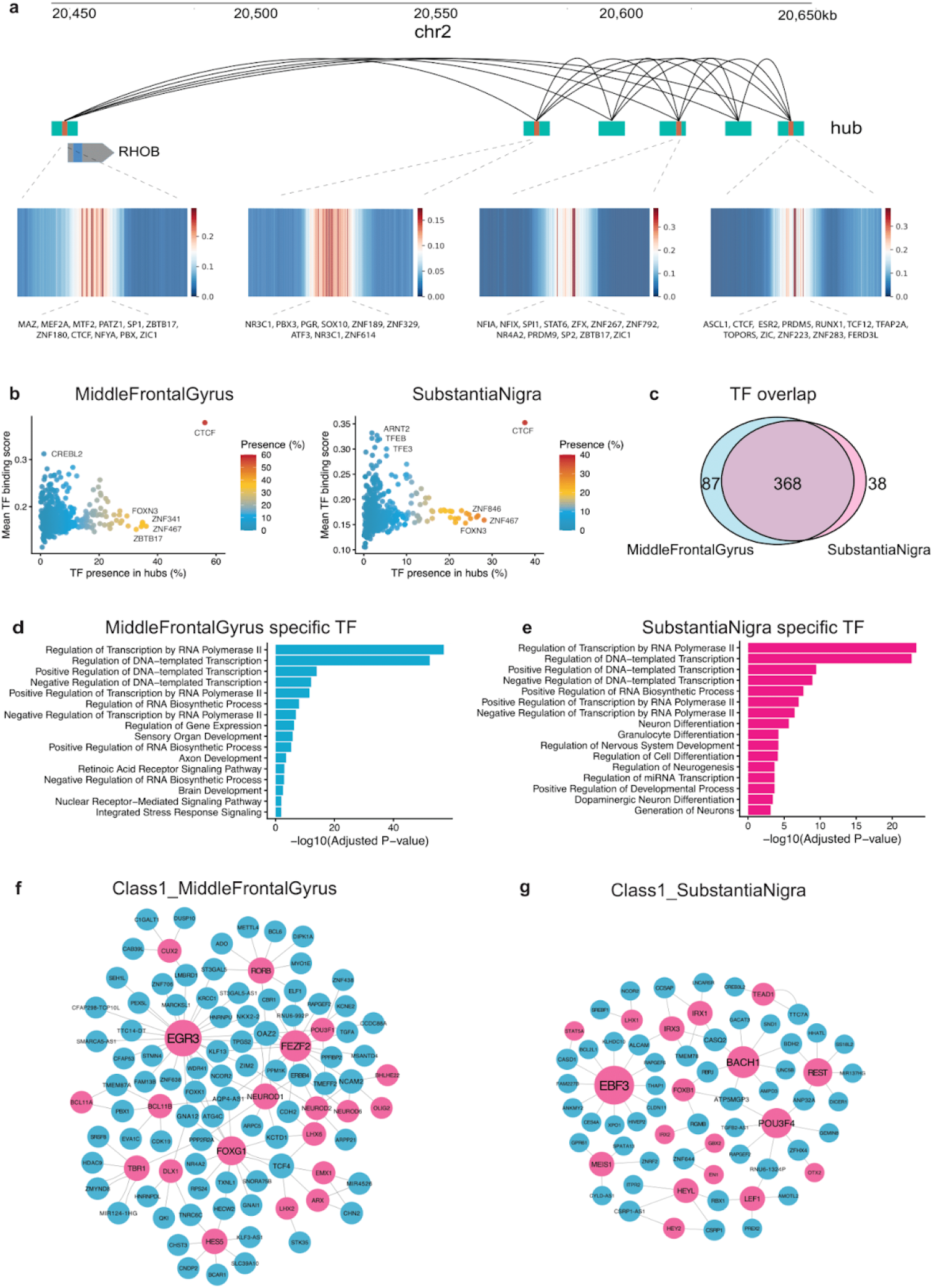
Transcription factor programs underlying hierarchical enhancer hub regulation. **a**, Example of a promoter-anchored multi-way enhancer hub in the *RHOB* locus identified from H3K27ac HiChIP data in the human brain caudate region. Arcs indicate inferred enhancer–promoter and coordinated enhancer interactions within the promoter-centered hub. Heatmaps show scPrinter-inferred transcription factor (TF) binding profiles across individual enhancer hub elements, illustrating coordinated TF occupancy within a higher-order regulatory assembly. **b**, TF prevalence versus mean binding score across enhancer hubs in middle frontal gyrus (left) and substantia nigra (right). Each point represents a TF, colored by the fraction of enhancer hubs in which binding is detected. Broadly acting and architectural TFs, including CTCF and ZNF family members, exhibit high prevalence and strong binding, defining a conserved transcriptional backbone. **c**, Overlap of TFs detected within enhancer hubs between middle frontal gyrus and substantia nigra. Numbers indicate TFs uniquely enriched in each region and those shared across both regions. **d**, Gene Ontology enrichment analysis of middle frontal gyrus–biased TFs, highlighting developmental and regulatory programs related to axon development, retinoic acid signaling, sensory organ development, and transcriptional regulation. **e**, Gene Ontology enrichment analysis of substantia nigra–biased TFs, showing enrichment for neuronal lineage specification, dopaminergic neuron differentiation, neurogenesis, and nervous system development. **f**, TF–gene regulatory network for region-restricted (Class 1) enhancer hubs in the middle frontal gyrus. Nodes represent TFs (pink) and target genes (blue), with edges indicating inferred regulatory connections mediated through enhancer hubs. **g**, TF–gene regulatory network for region-restricted (Class 1) enhancer hubs in the substantia nigra, illustrating distinct but structured TF–gene connectivity underlying region-specific regulatory programs.

Across the six brain regions analyzed, we identified approximately 300–500 TFs with detectable binding within enhancer hubs per region. Despite substantial regional differences in hub composition, a large subset of TFs (210 TFs) was consistently observed across all regions (Supplementary Fig. 4a), indicating the presence of a conserved transcriptional core associated with enhancer hub regulation. Gene Ontology enrichment analysis of these shared TFs revealed over-representation of fundamental transcriptional and neuronal regulatory processes, including RNA polymerase–associated transcription, neuronal differentiation, and central nervous system development (Supplementary Fig. 4b). These findings suggest that enhancer hubs are scaffolded by a broadly conserved, pan-neuronal TF repertoire rather than being specified solely by region-restricted TFs.

To assess TF prominence within enhancer hubs, we quantified both the prevalence of each TF across hubs and their aggregated binding scores inferred by scPrinter. This analysis revealed a compact set of highly recurrent and strongly bound TFs that consistently occupy enhancer hubs across brain regions (Fig. 5b; Supplementary Fig. 5). Broadly acting and architectural TF families, including CTCF, ZNF, SP, and SOX family members, exhibited both high hub-level prevalence and strong binding scores, consistent with established roles in chromatin organization and enhancer–promoter communication. Notably, while these TFs were broadly shared, individual TFs displayed substantial variation in prevalence and binding intensity across hubs, indicating selective deployment of common TFs rather than indiscriminate accumulation (Fig. 5b; Supplementary Fig. 5). Together, these observations define a conserved TF backbone underlying enhancer hub regulation.

In addition to this shared TF core, we observed subsets of TFs that exhibited region-biased enrichment within enhancer hubs (Supplementary Fig. 4a). Given the pronounced region-specific enhancer hub architectures identified in the middle frontal gyrus and substantia nigra, we focused on these two regions for further analysis. We identified 87 TFs preferentially enriched within middle frontal gyrus enhancer hubs and 38 TFs enriched within substantia nigra enhancer hubs (Fig. 5c).

Gene Ontology analysis of region-biased TFs revealed distinct functional emphases aligned with regional identity. In addition to transcriptional regulation pathways, middle frontal gyrus-biased TFs were enriched for developmental programs related to axon development, sensory organ development, retinoic acid receptor signaling, and brain development (Fig. 5d). In contrast, substantia nigra-biased TFs showed enrichment for neuronal lineage specification and maturation pathways, including dopaminergic neuron differentiation, generation of neurons, regulation of neurogenesis, and nervous system development, together with transcriptional regulatory processes (Fig. 5e).

Consistent with these functional enrichments, several region-biased TFs identified here correspond to well-established regulators of regional neuronal identity and development. In the middle frontal gyrus, biased TFs included canonical cortical regulators such as NEUROD2^21^, TBR1^22,23^, FEZF2^24,25^, and CUX2^26,27^, all of which play central roles in corticogenesis, neuronal differentiation, and the establishment of cortical projection neuron identity. In contrast, substantia nigra–biased TFs were enriched for regulators closely linked to midbrain and subcortical neuronal programs, including EN1^28,29^, REST^30^, and LHX1^31^, which have been implicated in dopaminergic neuron development, maturation, and long-term maintenance of subcortical circuitry. The recovery of these canonical, region-relevant TFs directly from enhancer hub–associated regulatory networks provides independent support for the biological specificity of the inferred TF programs.

To investigate how region-biased TFs contribute to enhancer hub–mediated gene regulation, we constructed TF–gene regulatory networks using highly enriched and well-established TFs of regional neuronal identity and development in each region. These networks revealed that region-biased TFs targeted genes associated with region-specific enhancer hubs (Class 1), providing a mechanistic link between region-biased TF deployment and region-restricted regulatory programs.

Notably, these TFs also exhibited connections to genes associated with pair-specific (Class 2) and multi-region shared (Class 3) enhancer hubs (Supplementary Fig. 4c). This observation indicates that region-biased TFs are not exclusively dedicated to a single enhancer hub class. Instead, they participate in a shared regulatory repertoire that can be engaged across multiple hierarchical hub contexts. These patterns suggest that hierarchical enhancer hub specification may not rely on class-specific TF identities alone, but may instead reflect context-dependent combinatorial assembly involving TF cofactors, chromatin state, and promoter competence.

Together, these analyses support a hierarchical view of transcription factor organization within multi-way enhancer hubs in the human brain. A conserved set of broadly acting TFs underlies shared regulatory programs across regions, while distinct subsets of region-biased TFs are associated with region-restricted enhancer hubs, reflecting regional identity and specialization. Within this framework, the coordinated involvement of shared and region-biased TFs, together with other context-specific factors, may contribute to circuit-level regulatory diversity and region-specific vulnerability, providing a potential mechanistic link between higher-order chromatin architecture, transcriptional regulation, and disease-associated genetic variation.

## Discussion

Understanding how multiple enhancers coordinate to regulate gene expression remains a central challenge in interpreting three-dimensional genome organization, particularly in complex tissues such as the human brain. Although higher-order chromatin interactions and enhancer clustering have been widely observed, the lack of a quantitative, promoter-anchored framework has limited the ability to define multi-way enhancer cooperation as a coherent regulatory unit or to systematically connect chromatin architecture with transcriptional output and disease-associated variation. In this study, we introduce a promoter-centric framework that formally resolves multi-way enhancer hubs as higher-order transcriptional regulatory units, integrates statistical inference with long-read physical validation, and enables cross-context comparison of regulatory architecture. By applying this framework to human brain chromatin interaction data, we reveal a hierarchical organization of enhancer hubs that links conserved neuronal regulatory backbones with circuit- and region-specific regulatory programs, and further demonstrate that this organization stratifies genetic risk and transcription factor programs across the brain. Together, these results establish multi-way enhancer hubs as a functionally meaningful layer of regulatory organization and provide a generalizable foundation for interpreting higher-order genome architecture in development and disease.

A central conceptual shift in our framework is the treatment of the promoter as a functional integrator of multi-way regulatory input. Previous approaches to identifying enhancer hubs from chromatin interaction data have largely relied on graph-centric or region-based abstractions that aggregate pairwise contacts into dense interaction clusters. While effective for describing highly connected genomic regions, such formulations do not explicitly define enhancer cooperation as a promoter-anchored regulatory unit and often conflate spatial proximity with functional coordination. By contrast, the promoter-centric framework introduced here reframes enhancer hub detection as a local, conditional problem centered on the promoter as the site of regulatory integration. Rather than clustering genomic regions based on global connectivity, our approach explicitly models enhancer–enhancer cooperation within promoter-anchored interaction neighborhoods, enabling enhancer hubs to be defined by synergistic co-occurrence rather than pairwise interaction frequency alone. The incorporation of quantitative interaction weighting, distance-stratified null modeling, and stability-based filtering further distinguishes this framework from prior heuristic formulations. Together, these features allow enhancer hubs to be resolved as statistically supported, physically validated, and functionally interpretable higher-order regulatory units, providing a unified methodological basis for comparative analysis across tissues, conditions, and regulatory states.

By resolving enhancer hubs as promoter-anchored regulatory units, our framework reveals a hierarchical organization of regulatory architecture that is not apparent from pairwise interaction analyses. Across human brain regions, multi-way enhancer hubs are structured into layered regulatory programs, comprising broadly shared hubs that define a conserved neuronal backbone, superimposed pair-specific hubs that capture circuit-level regulation, and region-restricted hubs that encode regional identity and selective vulnerability. This hierarchy is mirrored in the distribution of disease-associated genetic variation: broadly acting neuropsychiatric traits preferentially localize to multi-region shared hubs, whereas genetic risk for cognitive, psychiatric, metabolic, and neurodegenerative phenotypes becomes progressively concentrated within circuit- and region-specific hub classes. Together, these results indicate that non-coding disease-associated variants are not uniformly distributed across regulatory elements but are embedded within higher-order enhancer hub architecture, providing a structured framework that links three-dimensional genome organization, transcriptional regulation, and disease susceptibility in the human brain.

Several limitations of the present study should be considered. First, our analyses are based on bulk H3K27ac HiChIP and ATAC-seq data, which capture aggregate regulatory architectures across heterogeneous cell populations within each brain region. As a result, the inferred enhancer hubs represent composite regulatory units rather than cell-type-resolved assemblies. Although the reproducibility of hub structure across biological replicates and their concordance with long-read Pore-C validation support the physical and regulatory relevance of these hubs, future integration with single-cell chromatin interaction and accessibility technologies will be required to resolve cell-type–specific enhancer hub organization and heterogeneity. Second, the framework relies on existing promoter annotations to define regulatory anchor points. While promoter-centric resolution is essential for modeling enhancer cooperation as a functional regulatory unit, alternative transcription start sites or context-dependent promoter usage may not be fully captured. Incorporation of transcript-resolved or dynamic promoter definitions may further refine enhancer hub inference and enhance the mapping of regulatory input to transcriptional output.

Looking forward, the promoter-centric framework introduced here provides a generalizable foundation for interrogating higher-order regulatory architecture beyond the specific context of the human brain. As chromatin interaction and accessibility technologies continue to advance toward higher resolution and single-cell modalities, this framework can be readily extended to resolve cell-type-specific enhancer hub organization, developmental dynamics, and regulatory plasticity across physiological and pathological states. Integration with perturbative approaches, including CRISPR-based regulatory editing, will further enable systematic testing of enhancer hubs as functional units rather than isolated elements. More broadly, by defining enhancer cooperation as a promoter-anchored, statistically grounded regulatory entity, this work establishes a principled approach for connecting three-dimensional genome organization with transcriptional control and disease-associated variation across diverse biological systems.

## Method

### HiChIP data preprocessing and promoter-centric representation

H3K27ac HiChIP data were processed using HiC-Pro (v3.1.0)^32^ at 5 kb resolution. Read alignment, filtering, and contact matrix generation followed default HiC-Pro best practices. Statistically significant chromatin interactions were identified using HiCDCPlus (v1.14.0)^33^, which models distance-dependent background interaction frequencies and reports interaction-level statistical significance.

Promoters were defined as transcript start sites (TSSs) derived from GENCODE gene annotations^34^ (GRCh38 v46). Loop anchors overlapping promoter intervals were labeled as promoters (P), and all remaining anchors were labeled as enhancers (E). Only promoter–enhancer (PE) interactions were retained for downstream analyses. To avoid artifacts from extremely proximal contacts and poorly powered long-range interactions, only PE interactions with genomic distances between 10 kb and 2 Mb were considered for enhancer hub inference.

### Quantitative weighting of enhancer–promoter interactions

Each PE interaction was assigned a quantitative weight integrating interaction strength, statistical confidence, and distance dependency. Distance-dependent background behavior was modeled using fixed genomic bins:

[0,10 kb, 25 kb, 50 kb, 100 kb, 250 kb, 500 kb, 1 Mb, 2 Mb] Each interaction was assigned a bin.

We implemented 11 weighting schemes. Let *x* = counts, (*D*) distance, and (*q*) q-value:

1. count_only

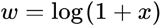

2. sig_only

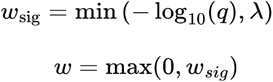

where λ is the significance cap (default λ=10) and missing significance is treated as 0.

3. distance_only (negative control; higher weight for proximal)

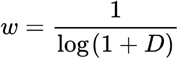

assigning higher weights to more proximal interactions.

4. count_sig

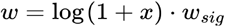

where *w*_*sig*_ if present, else 1.

5. count_sig_plus_dist_linear

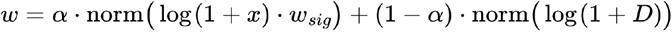

with α = 0.08 and *norm* denoting max-normalization to ([0,1]).

6. bin_percentile

Within each distance bin, counts were converted to empirical percentiles:

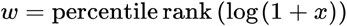

7. bin_percentile_plus_sig

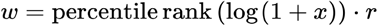

8. bin_log_ratio_sig

Within each distance bin, define expected count as the bin median:

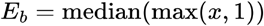

Compute log-ratio enrichment:

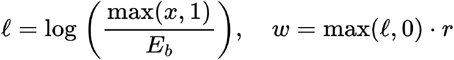

9. bin_diff_global

Within each distance bin, subtract the bin median in log-count space:

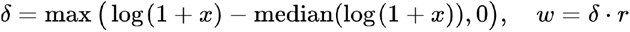

10. bin_diff_binmax

Compute (\delta δ) as above, then normalize within bin by bin maximum:

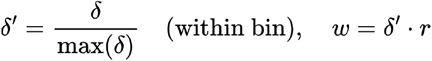

11. zscore_residual (requires HiCDCPlus outputs *mu, sdev*)

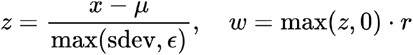

with ϵ = 10 ^−10^

To avoid overly aggressive multiplicative scaling by statistical significance, we defined a bounded reliability factor:

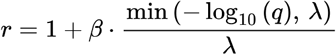

where λ is the significance cap (default λ=10) and β is a scaling factor (default β=0.5).

For the primary analyses, we used the distance-stratified log-enrichment weighting scheme (bin_log_ratio_sig). All other schemes were evaluated in sensitivity analyses to confirm robustness of hub detection.

### Promoter-conditioned enhancer co-membership networks

For each promoter *p*, we collected its associated enhancers {e_1_, … e_*n*_} from PE interactions, requiring a minimum enhancer degree of n ≥ 3,

We defined a promoter-conditioned enhancer–enhancer co-membership score that captures cooperative contribution conditional on the promoter. The co-membership score increases when two enhancers exhibit large promoter coupling relative to the aggregate enhancer weight landscape at that promoter. In the implemented log-space formulation, for enhancer and with nonnegative weights *w*_*i*_,*w*_*j*_, define:

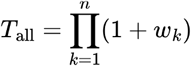

The (unnormalized) synergy-like score is:

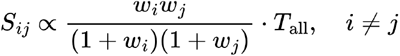

In practice, we compute in log space for numerical stability:

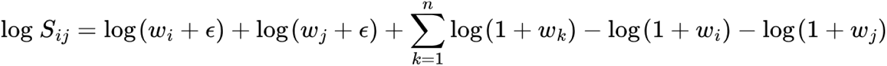

with ϵ = 10 ^−10^, and set diagonal entries to zero.

We further implemented co-membership normalization:

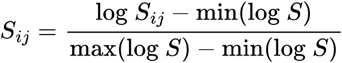

All normalized matrices were symmetrized, non-finite values were set to zero, and self-edges were removed.

### Hub calling by Leiden community detection and hub quantification

For each promoter, the normalized co-membership matrix defined an undirected weighted enhancer–enhancer graph. Enhancer hubs were identified using Leiden community detection with resolution parameter set to 1.0, allowing multiple hubs per promoter.

Hubs were ranked by dominance, defined as the sum of weighted node strengths within each community:

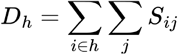

The highest-ranking hubs per promoter were retained as candidate hubs. For each hub *h* with *n*_*h*_ enhancers, we also computed:

Internal score:

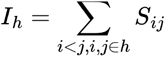

Weighted density:

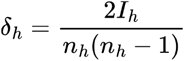

Graph density (QC metric): fraction of nonzero within-hub edges.

### Hub stability assessment

To assess robustness to sampling variability, we performed bootstrap resampling with 100 iterations. In each iteration *b*, interaction counts were resampled as:

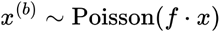

With *f =* 0.08, followed by full reweighting, co-membership construction, and Leiden clustering.

For each reference hub *A*, the most similar bootstrap hub *B* from the same promoter was identified using Jaccard similarity:

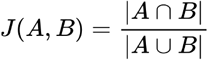

Hub stability was summarized by mean Jaccard similarity and stability rate, defined as the fraction of bootstrap iterations with *J* ≥ 0.5

### Distance-stratified null model and empirical significance

To distinguish structured enhancer cooperation from chance configurations while preserving distance dependency and interaction heterogeneity, we implemented a distance-stratified node-weight null model. For each distance bin, a global pool of PE weights was constructed.

For a given hub, null enhancer weight vectors were generated by resampling from the corresponding distance-bin pools (strict distance matching), followed by recomputation of the co-membership matrix and hub statistics. Empirical one-sided p-values were computed as:

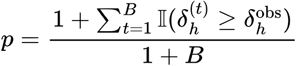

with B=1000 permutations, and adjusted across hubs using Benjamini–Hochberg FDR. Observed-to-expected ratios were calculated relative to the null median. Hubs with FDR < 0.05 and stability rate > 0.5 were considered high-confidence.

### Pore-C validation of enhancer hubs

Pore-C data were processed using the Pore-C-Snakemake pipeline (v0.3.0) ^18^. Reads were intersected with enhancer hubs to assess physical support for inferred multi-way interactions. A Pore-C read was considered supportive if it overlapped ≥3 hub elements, including enhancers and/or the associated promoter.

Validation metrics included the fraction of hubs supported by Pore-C reads, coverage of hub enhancers by Pore-C-supported bins, and the distribution of multi-contact order. Comparisons were performed across weighting strategies and hub confidence categories.

### Identification of replicable enhancer hubs across brain regions

For each brain region, enhancer hubs were identified independently in each replicate. To ensure robustness, we defined replicable enhancer hubs as candidate hubs identified at the same promoter in both replicates, sharing at least two enhancers, and significant in at least one replicate (FDR ≤ 0.05).

### Gene expression analysis

Hub-associated genes were defined as genes whose promoters participated in enhancer hubs. Gene expression levels were quantified using regional RNA-seq data and expressed as TPM. Sub-regions were merged within each of the six brain regions, while human individuals (A–D) were analyzed separately.

Expression distributions of hub-associated genes were compared with genes linked to pairwise enhancer interactions, randomly selected expressed genes, and all other expressed genes within each brain region.

### Cross-region sharing and hierarchical classification of hub-associated genes

Hub-associated genes were stratified by the number of brain regions in which they were observed. Pairwise overlap between regions was quantified using the Jaccard index. Pairwise overlap between regions i and j was quantified using the Jaccard index,

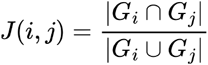

where and *G*_*i*_ and *G*_*j*_ denote the hub-associated gene sets for regions ii and jj, respectively. Jaccard indices were computed for all region pairs and visualized as a symmetric matrix heatmap. Genes were classified into multi-region shared (class 3), pair-specific (class 2), and region-specific (class 1) categories.

### Gene Ontology enrichment analysis

GO enrichment analyses were performed using clusterProfiler (v4.14.0)^35^ and Enrichr (v3.4)^36^. Analyses were conducted separately for hub-associated genes stratified by sharing class. Redundant terms were filtered using semantic similarity–based simplification implemented in simplifyEnrichment (v2.4.0) ^37^.

### GWAS variant enrichment analysis

To investigate the distribution of disease-associated genetic variation across higher-order regulatory architecture, we examined the overlap between enhancer hubs and genome-wide association study (GWAS) variants. GWAS data were obtained from the UCSC Genome Browser GWAS Catalog track^19^ (mirroring the NHGRI-EBI GWAS Catalog; accessed November 2025). The dataset was processed into BED format, with columns specifying genomic coordinates (chromosome, start, end) and the reported trait in the fourth column. This processing yielded 713,013 unique single-nucleotide polymorphisms (SNPs) linked to 31,944 distinct reported traits (as originally described in the source publications, prior to EFO ontology standardization). SNPs were intersected with enhancer hubs across brain regions. Enrichment analyses were performed at the enhancer-bin level using fisher test, stratified by hub class. Only associations with a p-value ≤ 0.01 and an odds ratio ≥ 5 were retained for further analysis.

### Transcription factor footprinting and network analysis

TF binding within enhancer hubs was inferred using scPrinter (v1.2.0)^20^ applied to matched ATAC-seq data across six brain regions. TF presence in enhancer hubs was defined as the fraction of hubs exhibiting a detectable footprint. Binding strength was quantified using aggregated scPrinter scores. Only TF associations with binding scores ≥ 0.1 and expression levels ≥ 5 TPM (from matched RNA-seq data) were retained for downstream analysis.

TF–gene regulatory networks were constructed by linking TF footprints within enhancer hubs to target genes via promoter-centered hub structure. Networks were visualized by Cytoscape (v3.10.3) ^38^ to illustrate combinatorial TF regulation across enhancer hub classes.

## Data visualization

The box plots, barplots, donut plots and radar chart were produced using ggplot2 (v.3.5.2; https://ggplot2.tidyverse.org), venn diagram were produced by VennDiagram (v1.7.3)^39^, heat maps were produced using pheatmap (v.1.0.13; https://cran.r-project.org/web/packages/pheatmap/index.html) in R (v.4.4.3) and UpSet plot were produced by ComplexUpset (v1.3.3)^40^. The representative tracks were produced using Gviz (v.1.50.0)^41^.

## Data availability

The GM12878 H3K27ac HiChIP data were obtained from the Gene Expression Omnibus (GEO) under accession number GSE188405. Pore-C data were downloaded from GEO under accession number GSE149117. H3K27ac HiChIP and ATAC-seq data from human brain regions were obtained from GEO under accession number GSE147672. Brain region RNA-seq data were also obtained from GEO under accession number GSE127898.

## Code availability

All code developed for multi-way enhancer hub inference and downstream analyses is publicly available on GitHub at https://github.com/YidanSunResearchLab/PEhub.git

## Acknowledgements

We thank Dr. Rich Head for insightful discussions and continuous support of this project. We thank all members of our department and institutes for fostering a collaborative and supportive research environment.

## Authors’ contributions

J.T. and Y.S. conceived the study. J.T. developed the computational framework, performed data analyses, and generated all figures. Y.S. supervised the project, guided methodological development, and contributed to data interpretation. Y.S. drafted the initial version of the manuscript, and J.T. and Y.S. jointly revised and finalized the manuscript.

## Ethics declarations

## Supplementary information

Supplementary Table 1: Replicable multi-way enhancer hubs identified across human brain regions.

**Supplementary Figure 1:**
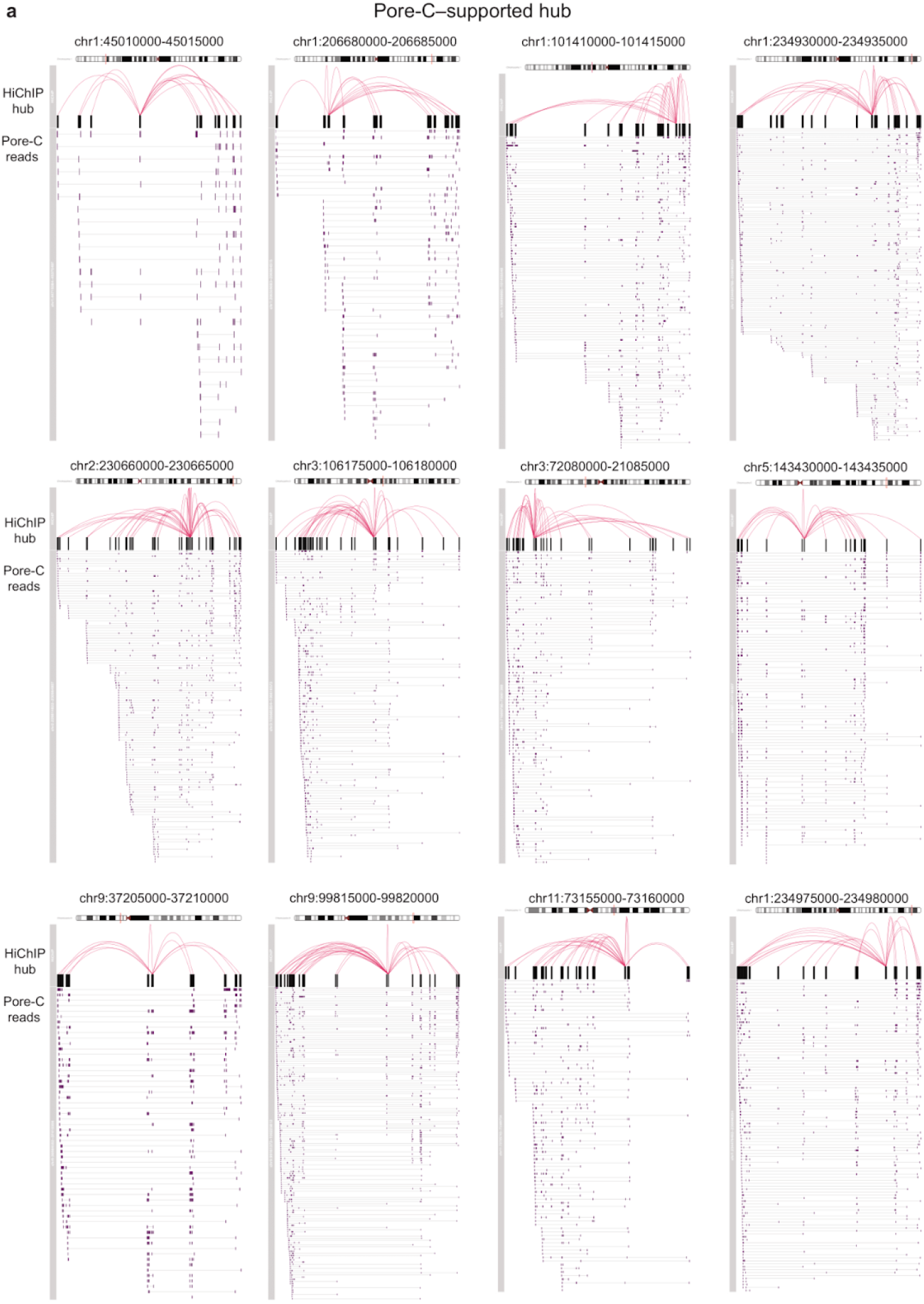
Examples of multi-way enhancer hubs supported by Pore-C reads. **a**, Representative examples of high-confidence multi-way enhancer hubs identified from GM12878 H3K27ac HiChIP data and independently supported by single-molecule Pore-C reads. A hub is considered supported if individual Pore-C reads overlap at least three hub elements, including enhancers and/or the associated promoter. For each locus, HiChIP-inferred enhancer hub structures (top) are shown together with aligned Pore-C multi-contact reads (bottom), illustrating direct physical evidence for higher-order chromatin assemblies.

**Supplementary Figure 2:**
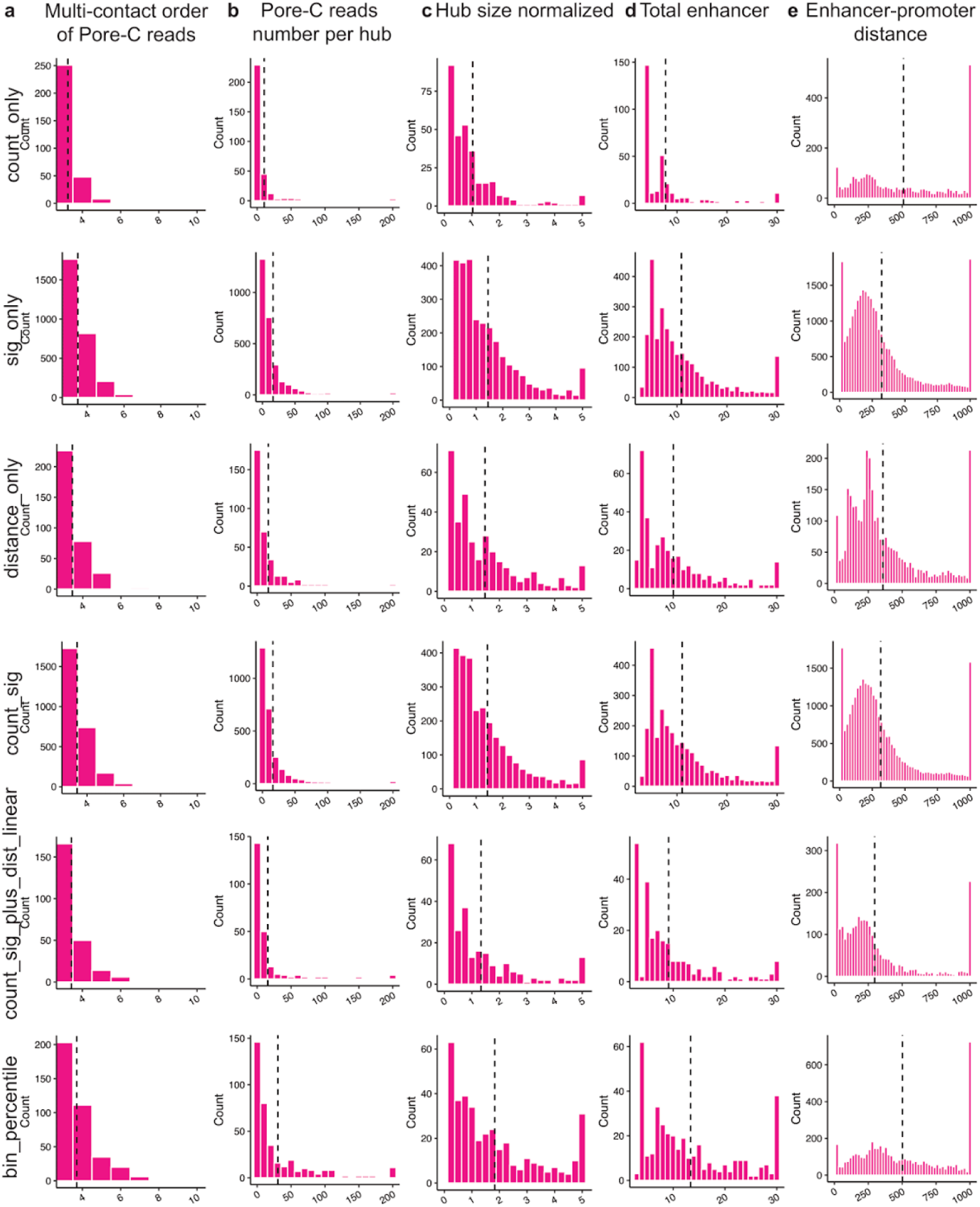
Pore-C validation of multi-way enhancer hubs across six alternative interaction weighting schemes. **a**, Distribution of multi-contact order (k) for Pore-C reads supporting enhancer hubs, reflecting higher-order chromatin interactions beyond pairwise contacts. High-confidence hubs are compared with filtered-out hubs. **b**, Distribution of the total number of supporting Pore-C reads per hub for high-confidence and filtered-out hubs. **c**, Supporting Pore-C reads per hub normalized by the number of enhancers within each hub, controlling for hub size effects. **d**, Distribution of hub size for high-confidence and filtered-out hubs. **e**, Genomic distances between enhancers and promoters within hubs, demonstrating that increased Pore-C support is not attributable to increased spatial proximity.

**Supplementary Figure 3:**
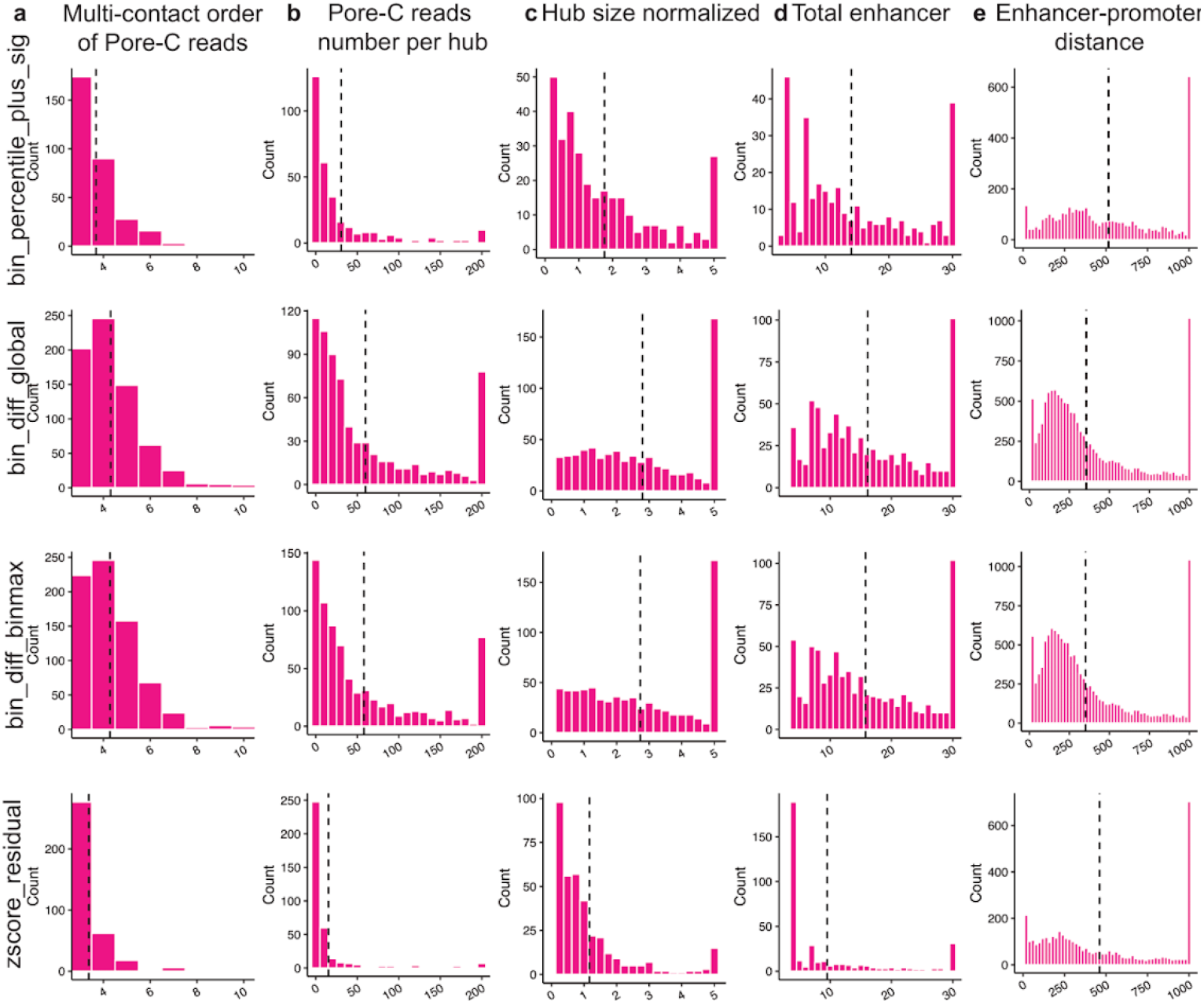
Pore-C validation of multi-way enhancer hubs across four alternative interaction weighting schemes. **a**, Distribution of multi-contact order (k) for Pore-C reads supporting enhancer hubs, reflecting higher-order chromatin interactions beyond pairwise contacts. High-confidence hubs are compared with filtered-out hubs. **b**, Distribution of the total number of supporting Pore-C reads per hub for high-confidence and filtered-out hubs. **c**, Supporting Pore-C reads per hub normalized by the number of enhancers within each hub, controlling for hub size effects. **d**, Distribution of hub size for high-confidence and filtered-out hubs. **e**, Genomic distances between enhancers and promoters within hubs, demonstrating that increased Pore-C support is not attributable to increased spatial proximity.

**Supplementary Figure 4:**
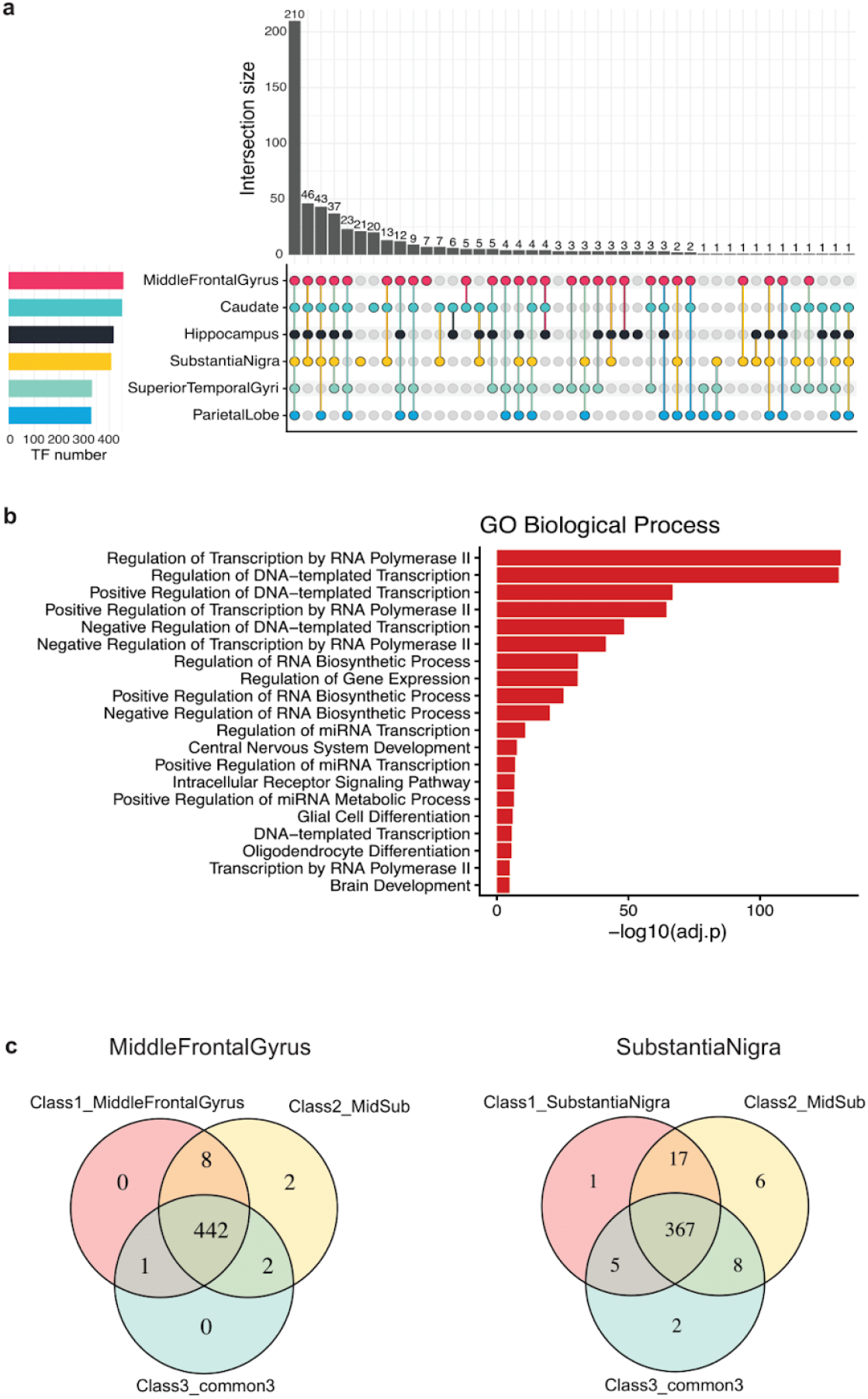
Transcription factor sharing and class-specific deployment across enhancer hub hierarchies. **a**, UpSet plot summarizing the overlap of transcription factors (TFs) detected within enhancer hubs across six human brain regions. Bars indicate the number of TFs present in the indicated combinations of brain regions, highlighting a large set of broadly shared TFs alongside region- or subset-specific TFs. The bar plot on the left shows the total number of TFs detected in enhancer hubs for each brain region. **b**, Gene Ontology biological process enrichment analysis of TFs shared across all six brain regions. Enriched terms highlight core transcriptional and neuronal regulatory processes, including RNA polymerase II–associated transcription, regulation of gene expression, and central nervous system development. Bars indicate −log10 adjusted *P* values. **c**, Venn diagrams showing the distribution of TFs across enhancer hub classes for middle frontal gyrus (left) and substantia nigra (right). TFs are stratified by association with region-restricted hubs (Class 1), pair-specific hubs shared between middle frontal gyrus and substantia nigra (Class 2), and multi-region shared hubs (Class 3). Numbers indicate TF counts in each intersection, illustrating that region-biased TFs participate across multiple hierarchical hub classes rather than being restricted to a single regulatory layer.

**Supplementary Figure 5:**
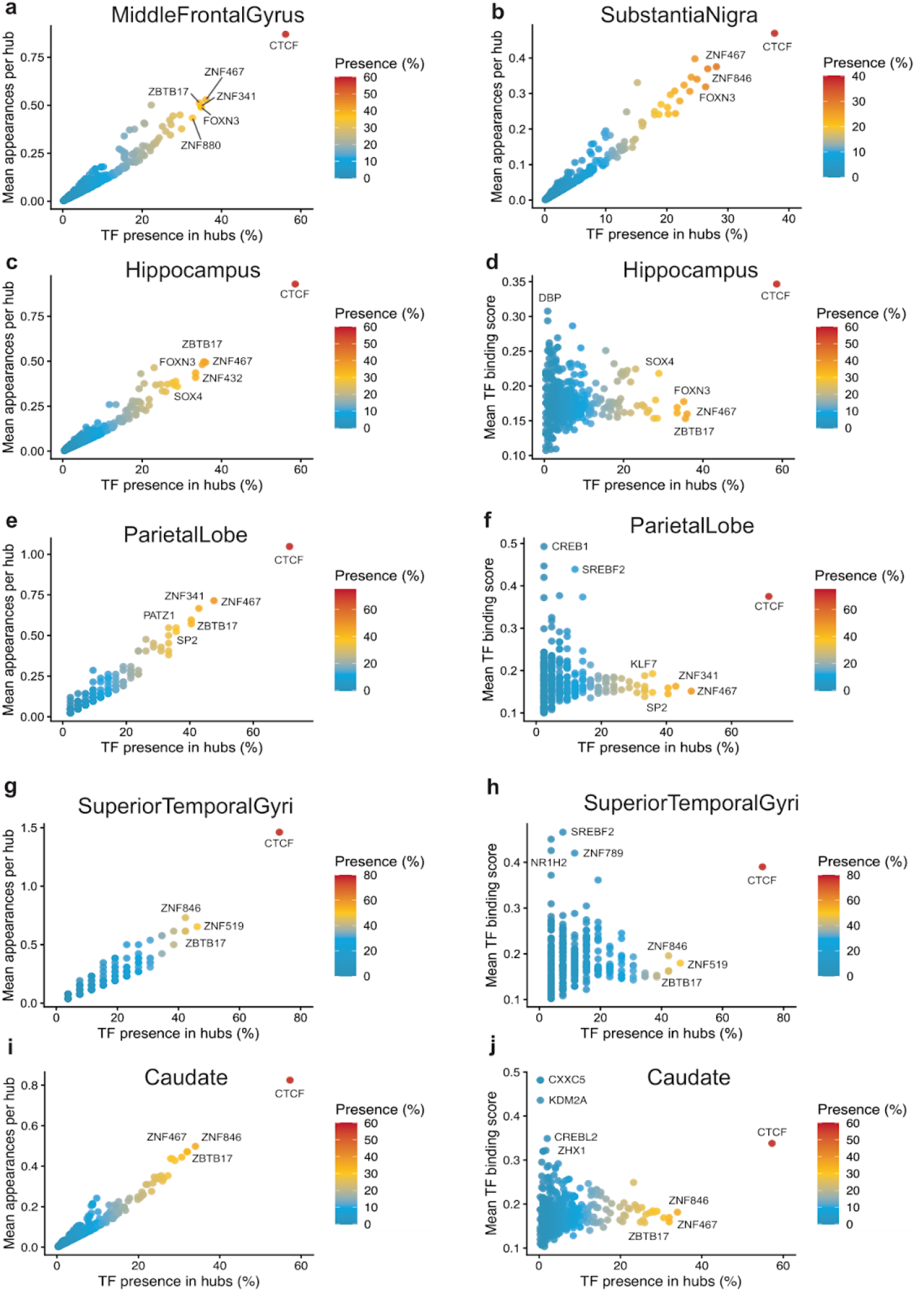
Transcription factor prevalence and binding strength across enhancer hubs in individual brain regions. **a–h**, Scatter plots summarizing transcription factor (TF) prevalence and binding characteristics within multi-way enhancer hubs across six human brain regions. For each region, TFs are plotted according to their presence in enhancer hubs (x-axis; percentage of hubs containing at least one footprint for the TF) and either their mean appearances per hub (left column) or mean scPrinter TF binding score (right column), reflecting binding frequency and binding strength, respectively. Panels correspond to individual brain regions as follows: **a**,**b**, middle frontal gyrus; **c**,**d**, hippocampus; **e**,**f**, parietal lobe; **g**,**h**, superior temporal gyrus; **i**,**j**, caudate. Each point represents a single TF detected by scPrinter footprinting in region-matched ATAC-seq data and RNAseq data (binding scores ≥ 0.1 and expression levels ≥ 5 TPM). Points are colored by TF presence across hubs (color scale), highlighting TFs that are broadly recurrent versus sparsely deployed. Selected TFs with high prevalence and/or strong binding are labeled to illustrate representative architectural and regulatory factors, including broadly acting TFs (for example, CTCF, ZNF family members, SP family members) and region-biased factors. Together, these analyses illustrate that enhancer hubs within each brain region are characterized by a compact set of highly recurrent, strongly bound TFs embedded within a broader distribution of lower-prevalence factors, supporting the existence of a conserved transcription factor backbone with region-specific modulation.

